# Quantitative traits under selection: derivations of distributions, higher moments, and the effects of recombination

**DOI:** 10.1101/2021.07.30.454547

**Authors:** Reginald D. Smith

## Abstract

The mathematical theory of quantitative traits is over one hundred years old but it is still a fertile area for research and analysis. However, the effects of selection on a quantitative trait, while well understood for the effects on the mean and variance, have traditionally been difficult to attack from the perspective of analyzing the probability density of the breeding values and deriving higher (third and fourth) moments as well as analyzing the impact of recombination. In this paper, the exact formula for the breeding value distribution after selection is derived and, using new integral tables, the first four moments are given exact expressions for the first time. In addition, the effects of recombination on the full distribution of breeding values are demonstrated. Finally, the changes of GXE covariance in the selected parent population caused by factors similar to the Bulmer Effect are also investigated in detail.

**Mathematics Subject Classification (2020):** MSC code 92D10 · MSC code 92D15

## 1 Introduction

A hallmark in the development of genetics, the theoretical underpinnings of quantitative genetics by Fisher [Fisher, 1918] was brought about by the at first seemingly irreconcilable facts of the inheritance of traits demonstrated by Mendelian laws of inheritance at a single locus and the correlations between relatives of traits demonstrated by biometrics. The complete description of the relationship between phenotype, genetics, and environment remains complex and evolving but in modern parlance the basic description of the relationship between random variables *P, G*, and *E* representing the phenotypic values, breeding values, and environmental values is given by

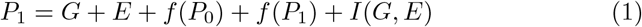

Here the phenotype in progeny of generation 1 is given by the sum of breeding and environmental values as well as forms of non-genetic inheritance and the gene-environment interaction *I*(*G, E*). Non-genetic inheritance can take forms as vertical inheritance from the parent generation *f* (*P*_0_) such as maternal inheritance [Kirkpatrick & Lande, 1989] or oblique cultural inheritance from non-parents in the previous generation [Cavalli-Sforza & Feldman, 1981]. It can additionally include horizontal inheritance from other members of the same generation, *f* (*P*_1_) such as horizontal cultural inheritance. In the simplest model, and the only one we will address, this relationship is simplified to

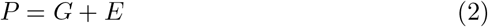

The term, *G*, is typically designated the breeding value and is based on the average phenotypic value of progeny that can be expected by mating an individual with another random individual in the population. The expected progeny phenotype given midparent phenotype is described by the Breeder’s Equation [Lush, 1943, Lush, 1948]

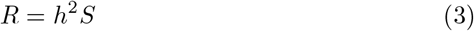

*S*, is the difference between the average phenotypic value of the parents and the mean of the population and *R* is the difference between the phenotypic value of the offspring and the mean of the population. The variable *h*^2^ is the narrow heritability representing the proportion of total population variance that is associated with the variance of additive breeding values in the population. Where the average environmental effect is assumed to be zero, randomly mating an individual of breeding value *G* is expected to give *R* = *h*^2^*G/*2. Historically and even with the advent of the use of molecular genetics in animal breeding, breeding values are defined by the expected progeny values from individuals rather than expected individual phenotypes.

The Breeder’s Equation is the earliest and most succinct relationship describing the effects of phenotypic selection from the parent population on the progeny population. Further work on this topic, in particular addressing the moments of the progeny generation was continued by Bulmer [Bulmer, 1971, Bulmer, 1974, Bulmer, 1985] who demonstrated the Bulmer Effect that showed changes in variance due to phenotypic selection were due to changes in linkage disequilibrium which decayed over time. Analyzing the higher moments were often difficult due to the distortion of normality induced by selection. Selection not only changes the variance but also skew, excess kurtosis, etc. due to higher order (three, four loci) linkage disequilibrium. The non-normal nature of the distribution made recourses to various techniques such as Gram-Charlier series [Bulmer, 1985, Zeng, 1987] and the method of cumulants [Bürger, 1993, Barton & Turelli, 1990, Barton & Turelli, 1991, Barton & Turelli, 1994] most effective for calculating the moments and modeling the effects of recombination.

Until recently, definite integrals were not known for most calculations on these equations since they necessarily contained the expression for the cumulative distribution function of the normal distribution, often called the error function (erf). However, given several sources [Ng & Geller, 1969, Ng & Geller, 1971, Korotkov & Korotkov, 2020], especially the recent translated work of Korotkov, the definite integrals necessary to derive the exact form of the breeding value distribution under most conditions, as well as its moments, are now available.

In the first part of the paper we will derive the exact expression for the breeding value distribution subject to phenotypic selection and then re-derive the results for the first two moments using tables of integrals in Appendix A. While the results here are already widely known, this method will allow the investigation of these moments to demonstrate not only are their results valid for arbitrary phenotypic selection schemes or the strength of selection, but also to provide exact equations to calculate the values of these moments for interval, truncated directional, stabilizing, and disruptive selection schemes using only the values of the phenotypic selection thresholds. This allows the values of the post-selection phenotypic mean and the change in phenotypic variance to be easily calculated directly. Following this we will show how these results can be easily applied to situations of GXE covariance.

In the following section, new results will describe the distribution of the environmental values of the parent generation after within-generation phenotypic selection. The moments of this distribution accentuate the concept that phenotypic selection selects not just breeding values but also environmental values as well. This non-random selection of breeding and environmental values by phenotypic selection induces a GXE covariance between breeding and environmental values and also a change in the phenotype-breeding value covariance in the parent generation post-selection. These do not change the expected values of progeny moments but can be used to investigate the relationship between traits and environment in a population selected by phenotype.

Subsequently, the between generation changes in the breeding value distribution due to recombination will be defined. The applicability is limited to the two locus linkage disequilibrium affecting the variance described by the Bulmer Effect, but will allow an exact description of the change in breeding values between the within-generation selected parents and their offspring.

Finally, the third and fourth moments of the breeding value distribution post-selection will also be introduced and we will confirm the results of [Barton & Turelli, 1994] that indicate the skew (third moment) of the breeding value does not show large distortion, even under strong selection, but the excess kurtosis (fourth moment) can show large changes under disruptive selection.

## 2 The Breeding Value Distributions and Phenotypic Selection

Our starting point will be equation 2. In the first model, we investigate an infinite population where the breeding and environmental values are considered statistically independent. The mean of the phenotypic, breeding value, and environmental value distributions are assumed to be zero for simplicity so non-zero values of phenotype represent *S* or *R* from the Breeder’s Equation. The original population phenotypic variance is given by *σ*^2^. The breeding values represent additive genetic variance without additive epistasis and have variance *h*^2^*σ*^2^ while the environmental values have variance (1 − *h*^2^)*σ*^2^. The initial phenotype marginal distribution is normally distributed as given by

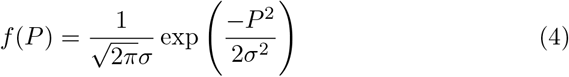

The marginal distributions for the breeding values and environment are both normal distributions given by

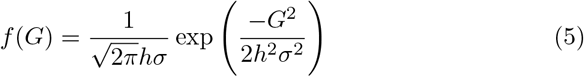

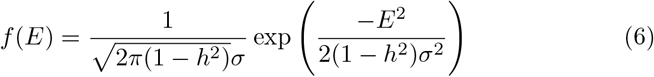

The joint distribution of the breeding values, *f* (*G, E*) absent GXE covariance is given by their product

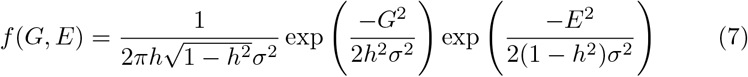

Given *E* = *P* − *G*, equation 7 can be rewritten as

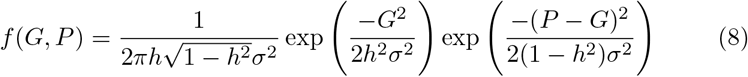

The equation 8 will be our starting point in investigating the effects of phenotypic selection on the breeding value distribution. For the interval selection of phenotypes between a lower threshold *T*_*L*_ and upper threshold *T*_*H*_, *T*_*H*_ *≥ P ≥ T*_*L*_, we must first integrate *P* between these values

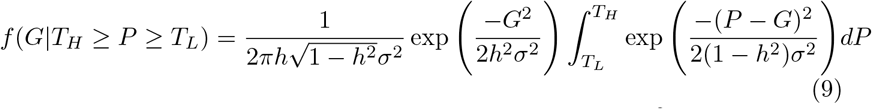

The evaluations of integrals of the form ∫exp (− (*ax* + *b*)^2^)*dx* formally derives the cdf of the normal distribution, *Φ*(*ax* + *b*) which is often explicitly defined as the error function 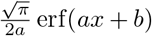. To solve these otherwise difficult expression, various integrals, with their sources, are listed in Appendix A. To evaluate equation 9 we use integral 49 to derive

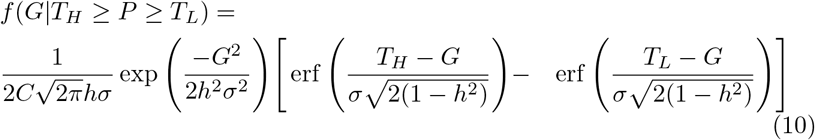

In the expression above *C* is a constant represented by the sum over all values of *G* in order to normalize the conditional pdf

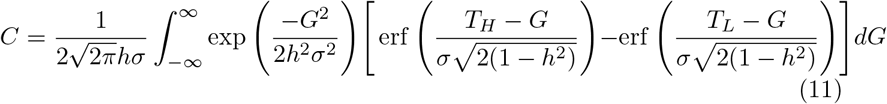

Equation 11 can be evaluated using two separate integrals and equation 51 to find

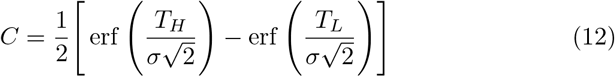

The final equation for the distribution of breeding values of interval selection gives

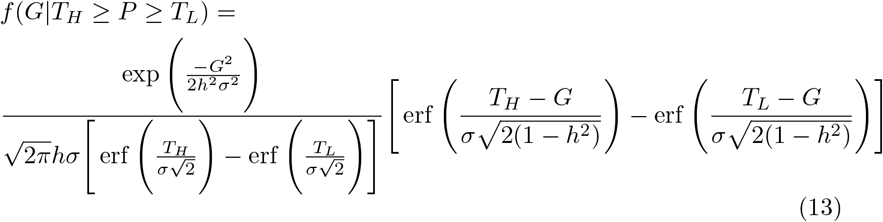

Equation 13 is the conditional distribution of breeding values subject to interval phenotypic selection between two phenotypic values. For stabilizing selection around the mean, *T*_*H*_ = *T* and *T*_*L*_ = −*T*.

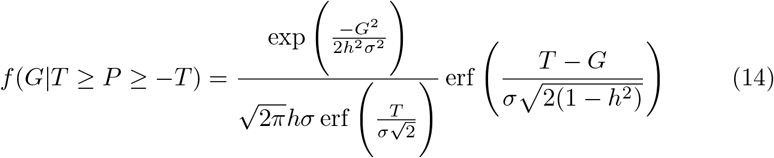

For truncated directional selection above a threshold *T*, it can easily be modified by setting *T*_*H*_ = *∞* and *T*_*L*_ = *T* while noting erf *∞* = 1 and erf 0 = 0.

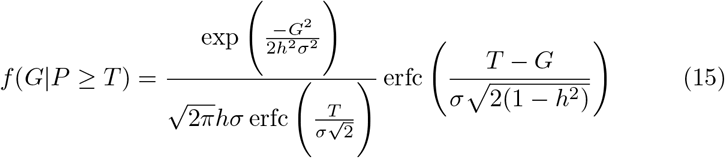

The new expression above, erfc(*x*), is the complimentary error function defined as erfc(*x*) = 1 − erf(*x*). In a similar fashion, disruptive selection can be analyzed by reincorporation *T*_*L*_ and *T*_*H*_ in separate integrals with integral limits bounding infinity on both sides and accounting for a normalization constant of twice the size as in truncated directional selection.

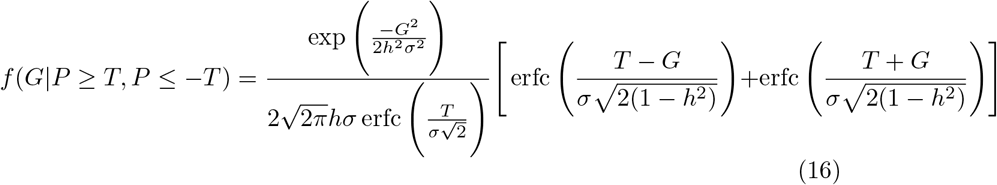

In all cases, the threshold phenotypic values can also be calculated based on percentages. For example for choosing the top *p* proportion of organisms for selection, *T* = *Φ*^−1^(1 − *p*) where *Φ*^−1^ is the quantile function of the phenotypic normal distribution.

Equations 13 and 15 are representative of similar forms discussed in [Slatkin, 1970, Karlin, 1979, Barton & Turelli, 1994]. Using notations from these papers, the breeding values are described as

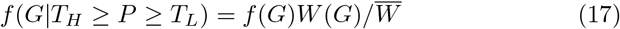

*W* (*G*) and 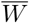 are the fitness function and the average fitness respectively. Therefore, the numerator erf and erfc functions of *G* are the fitness functions and the denominator erf or erfc constants are the average fitness.

## 3 The first two moments of breeding values subject to phenotypic selection

Now that we have described the transformation of the breeding value distribution by within generation selection we can now calculate the moments analytically using the equations in Appendix A. First using equation 52. The results for interval and truncated directional selection are

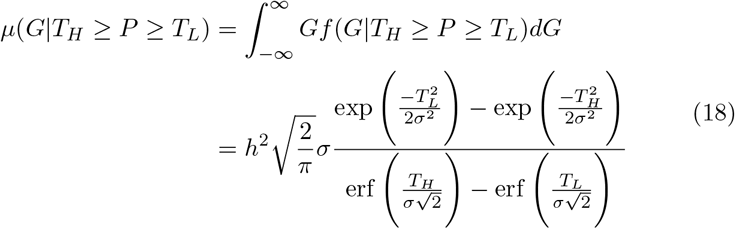

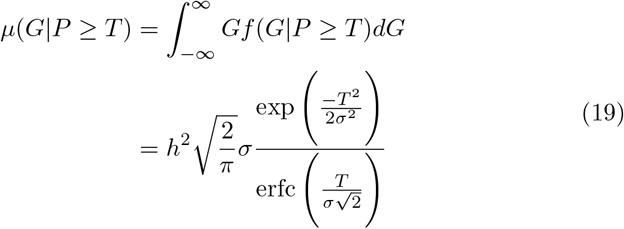

As expected the means of stabilizing and disruptive selection are zero. The complexity of these expressions can be simplified by looking at the mean of the truncated phenotype distribution as derived below for interval and truncated directional selection.

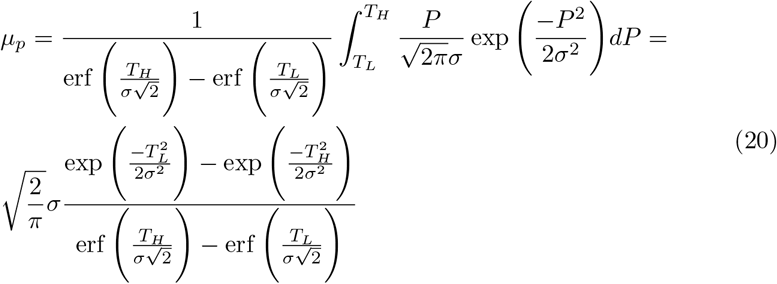

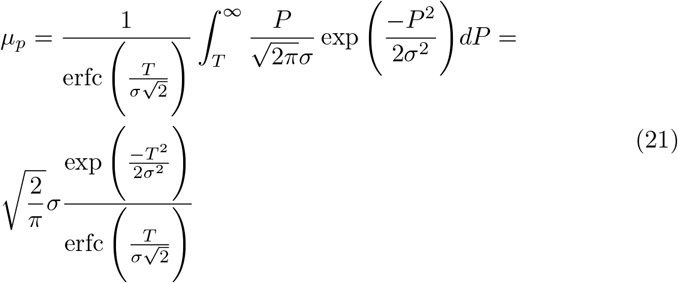

Therefore, it is clear that the derived expressions in equations 18 and 19 are equivalent to the Breeder’s Equation, *h*^2^*µ*_*p*_.

Similarly, we can calculate the variances with equation 53, however, instead of calculating the moment around zero, such as to find the new mean breeding value, we must calculate the moment with the reference to the new mean breeding value, *µ*_*G*_ = *h*^2^*µ*_*p*_, to accurately determine the variance of the new distribution. For interval selection this becomes:

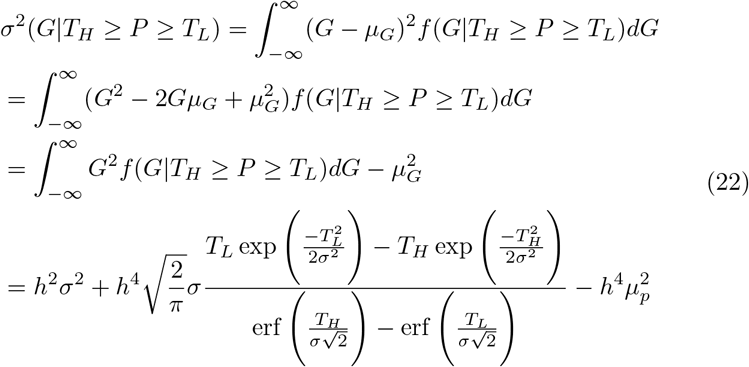

For truncated directional selection:

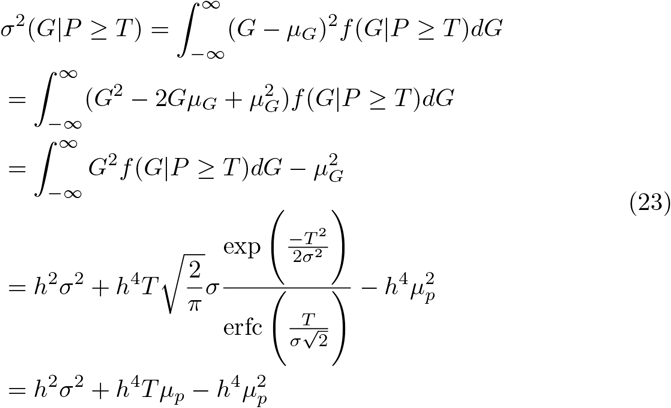

Once again, calculating the phenotypic variance around the new phenotypic mean makes it clear that this is the Bulmer effect. For interval selection and where *µ*_*p*_ is the phenotypic mean after within generation selection.

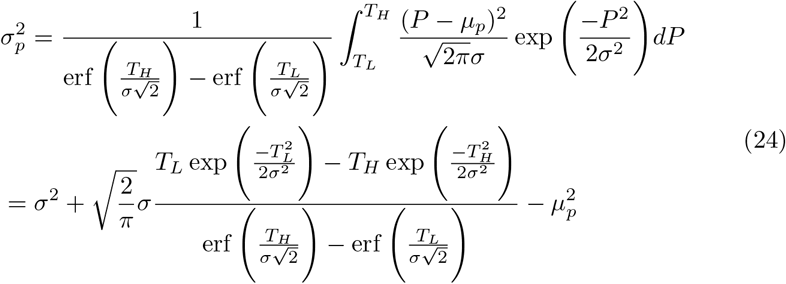

and for truncated directional selection

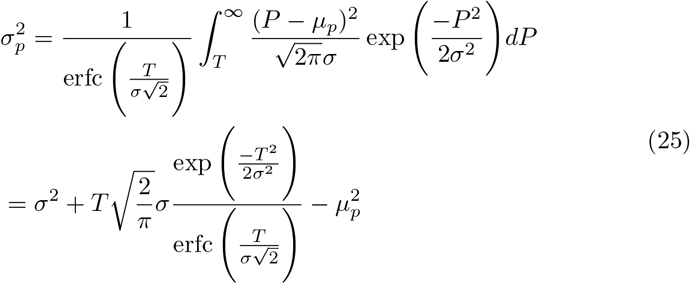

It is simple to see that the change in the second moment for the breeding value distribution is *h*^4^*Δσ*^2^ where *Δσ*^2^ is the difference between the original phenotypic variance, *σ*^2^ and the phenotypic variance after phenotypic selection. This is identical to the typical expression for the Bulmer Effect with the exception of a factor of 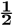. This variance, being the variance of the phenotype selected parents, has not yet incorporated the effects of recombination which will reduce *h*^4^ to *h*^4^*/*2 in the progeny generation.

### 3.1 Disruptive phenotypic selection

The disruptive selection variance is similar to the directional selection variance with the exception that the breeding value variance before selection is doubled and the final term subtracting the mean squared disappears so the change in additive variance is positive.

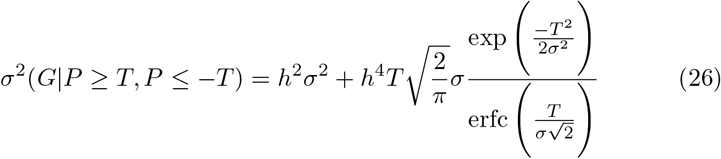

## 4 Gene x Environment Covariance

The breeding value distribution taking genotype x environment (GXE) covariance into account can be easily derived by modifying equation 7 to incorporate the correlation *r*_*GXE*_ and expressing the genotype variance before selection as 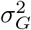 without referencing narrow heritability. The bivariate normal distribution of *G* and *E* is thus

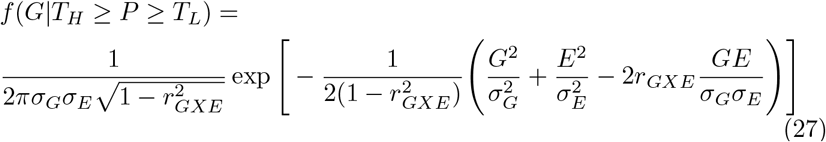

The equation for interval selection then becomes

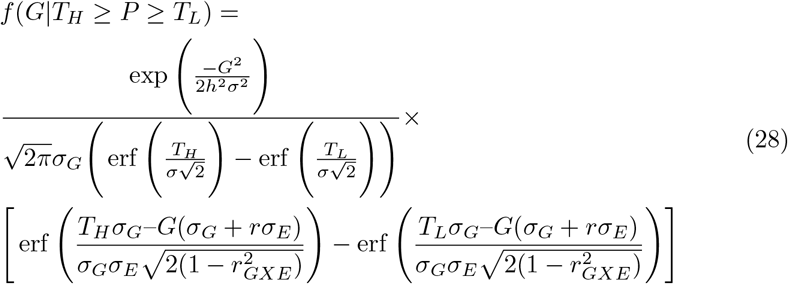

Similar to results in [Van Tienderen & De Jong, 1994, Mulder & Bijma, 2005, Bijma, 2020] the GXE covariance *σ*_*GXE*_ = *r*_*GXE*_*σ*_*G*_*σ*_*E*_ plays a crucial role in expected response. The first moment is now

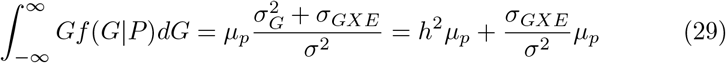

Similar with the variance

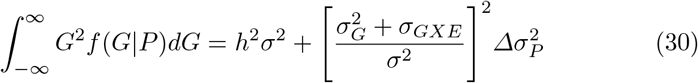

## 5 Environmental value distributions after within generation selection

For the selected phenotypes within the parent generation, we can calculate the distribution of environmental values similar to the breeding values. The joint distribution of *P* and *E* is given by

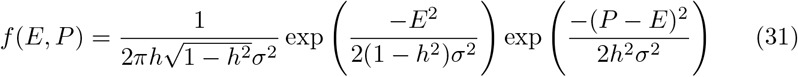

The distribution following within generation selection is then

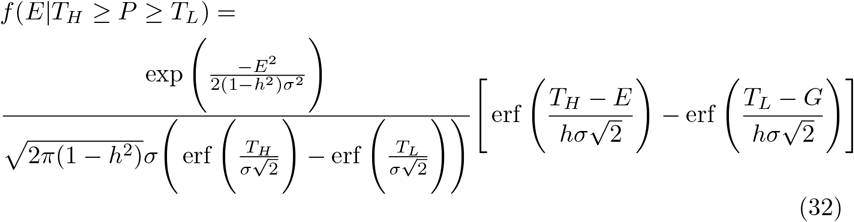

The first two moments follow as before and are given by

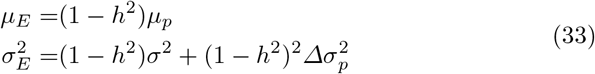

These demonstrate again how selecting based on phenotype does not randomly select environments. For any given single breeding value, if there is no *GXE* covariance then it encounters the full distribution of environments but selecting on phenotypes there is a non-random selection of environmental influences similar to the non-random selection of breeding values.

## 6 Beyond the Bulmer Effect: Other consequences of changes in variance

From the preceding section it was shown that environmental values are selected similar to breeding values as a result of within generation phenotypic selection. While this is widely expected, what is likely less understood is that phenotypic selection induces a covariance between the breeding values and environmental values even if *GXE* covariance was absent in the pre-selection population. Assuming no *GXE* in the original population, after phenotypic selection the variances are related by

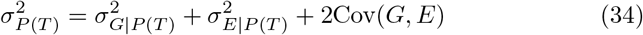

or

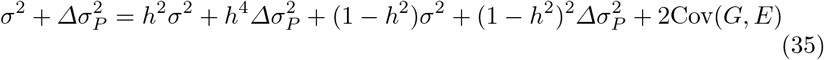

The final result is

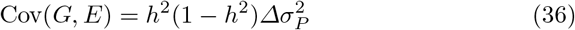

This shows that even if *GXE* covariance is not present in the original population pre-selection, *GXE* covariance is induced by any phenotypic selection that changes the phenotypic variance of the population. The magnitude of the covariance is directly proportional to the magnitude of the variance change and the product of the narrow heritability and environmental proportion of variance. This effect has a form and magnitude similar to the linkage disequilibrium covariance caused by the Bulmer Effect.

For any given change in variance the induced covariance effect is greatest when *h*^2^ = 0.5 so that *h*^2^(1 − *h*^2^) = 0.25. However, even for large or small values of *h*^2^ such as *h*^2^ = 0.2 or *h*^2^ = 0.8 the value of *h*^2^(1 − *h*^2^) = 0.16 so the phenotypic variance change is the most important factor in determining the final magnitude of the covariance.

For selection that reduces the variance such as interval, stabilizing, or directional selection, there is negative covariance between breeding and environmental values so that smaller breeding value individuals most likely have larger direct environmental influences on phenotype and vice versa. While there can be a heritable component to environmental variance [Zhang & Hill, 2005], environmental values are usually not directly inherited so this covariance is severely weakened or absent in the progeny population.

Similarly, the covariance between the phenotype and the breeding values, which is the breeding value variance, *h*^2^*σ*^2^, in the pre-selection population undergoes a change that is also related to the change in phenotypic variance from selection. Given *P* + *G* = 2*G* + *E* we can state the first relation

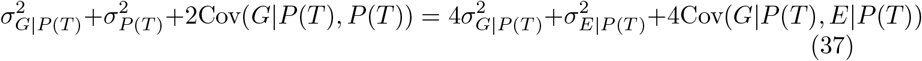

Which using the variance definitions from equations 35 and 36 gives us the final expression for covariance between breeding value and phenotype after within generation selection

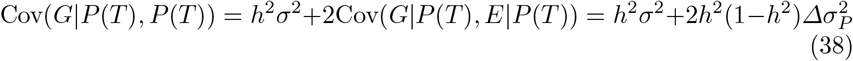

When there is no change in phenotypic variance, the phenotype and breeding value covariance reverts to its well-known value of *h*^2^*σ*^2^. Interestingly, amongst the population of selected parents from stabilizing or directional selection, where the additive variance is reduced, the phenotype and breeding value covariance declines. Since 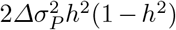 is not identical to the change in phenotypic variance due to sum of the differences due to the selected breeding and environmental values 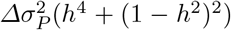, this is a change in both the covariance and correlation between phenotype and breeding value in the selected population making phenotype a less accurate correlate to breeding value in the selected parent population.

### 6.1 Bulmer Effect for correlated traits

A final derivation will show that the Bulmer Effect for correlated traits is relatively weak, implying that linkage disequilibrium caused in a trait correlated to the trait under selection is comparatively small except under circumstances of high correlation squared. The change in phenotypic variance of trait 1, 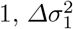 given the change in the phenotypic variance of trait 2, 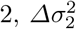, which is undergoing phenotypic selection, is given by

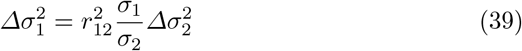

Thus the correlated Bulmer Effect is given by

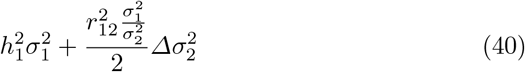

Given 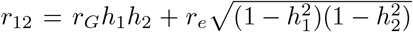, where *r*_*G*_ is the genetic correlation of additive trait loci and *r*_*e*_ the correlation of relevant environmental variables, if there is no environmental correlation between the traits (usually a strong assumption for closely correlated traits) and assuming the variances are normalized to be equal, the correlated Bulmer Effect is

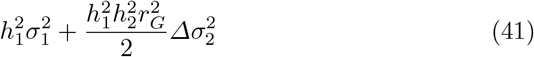

## 7 The effect of recombination on the breeding value distribution

To understand the full effect of recombination on the progeny breeding value distribution we will introduce an additive expression to the within generation selected breeding values that fully accommodates the effect of recombination on the variance and partially for higher moments. For a within generation breeding value distribution subject to phenotypic selection, *f* (*G*_1_) = *f* (*G*|*P*), the progeny breeding value distribution, *f* (*G*_2_|*G*_1_) becomes

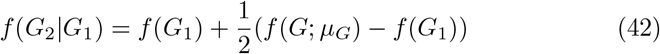

In this equation, *f* (*G*; *µ*_*G*_) is the normal distribution with mean *µ*_*G*_ = *h*^2^*µ*_*p*_ and variance *h*^2^*σ*^2^. So the difference between the normal distribution with the new mean and the starting additive genetic variance and the within selection breeding value distribution determines the change in the progeny breeding values. Note that since both *f* (*G*; *µ*_*G*_) and *f* (*G*_1_) sum to one over their entire range, there is no need to add another normalization constant for *f* (*G*_2_ | *G*_1_) and the first moments of both are equal and cancel out so that the mean breeding value of the selected parents and progeny remain the same. The only effects come with the higher moments that see the moments attributable to linkage disequilibrium reduced by one-half due to recombination thus recovering the traditional progeny variance expected by the Bulmer Effect. If variance is reduced by selection such as in stabilizing or directional selection, the larger variance of *f* (*G*; *µ*_*G*_) increases the variance between generations whereas if variance increases such as in disruptive selection, the variance is decreased between generations. Per [Barton & Turelli, 1994] higher moments are reduced even more so that the third moment is reduced to 1/4 of its value in the post-selection parent generation and the fourth moment is reduced to 1/8 of its value. Therefore, the expression above only partially accounts for the breakdowns of three, four, etc. loci linkage disequilibrium.

The effect of recombination on the breeding value distribution between generations can be informative as well as how recombination affects the frequencies of breeding values. As shown in Figure 1, that shows an example plot of 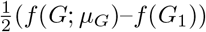 post truncated directional selection. The largest effect on breeding value frequencies due to recombination is at the mean value of the population where the frequency is reduced between generations if the variance was reduced by selection and vice versa where variance is increased by selection. This effect falls off rapidly though farther from the mean and actually reverses for intermediate values on both sides of the mean. Values far from the mean see almost no effect on their frequencies due to recombination.

**Fig. 1.**
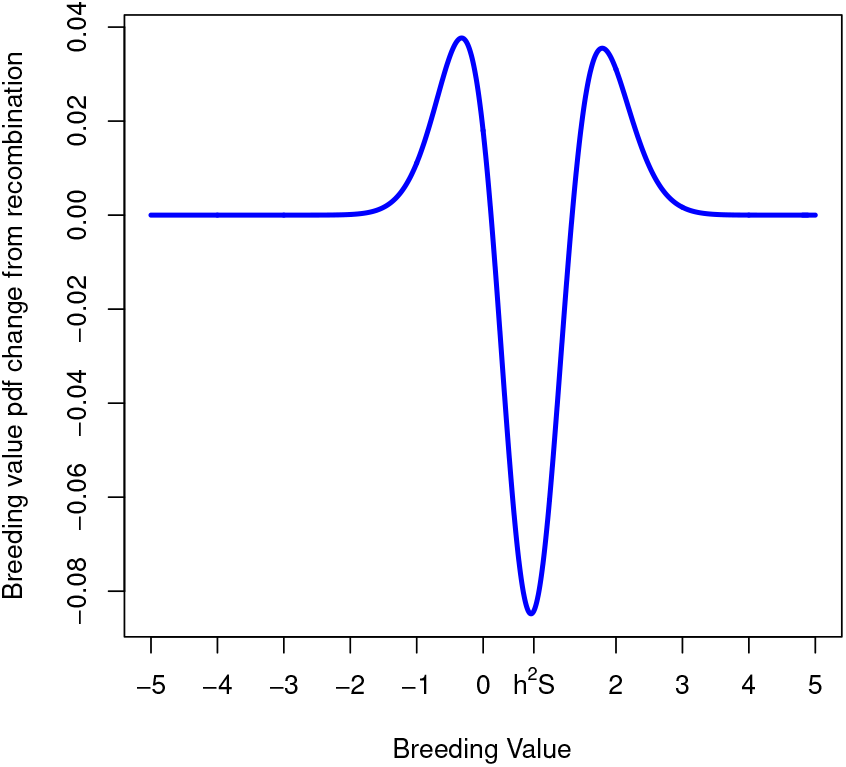
The change in the breeding value distribution induced by recombination after positive truncated directional selection in the parent generation. Created in R.

## 8 Higher moments

The expressions for skew (third moment) and excess kurtosis (fourth moment) are similarly calculated. To maintain brevity, these expressions will leave out most intermediate calculations to obtain final solutions.

### 8.1 Skew

The question of the importance of the third moment in quantitative genetics goes back to its earliest days when [Fisher et. al., 1932] investigated effects on the third moment for additive quantitative traits with few trait loci or traits depending on dominance effects. Skew was further investigated by [Barton & Turelli, 1994] and found to arise for third order (three loci) linkage disequilibrium and was not large for even strong selection. We will explore this in the derivations here.

For skew given the same assumptions for *µ*_*G*_ and *µ*_*p*_, first we calculate the third moment

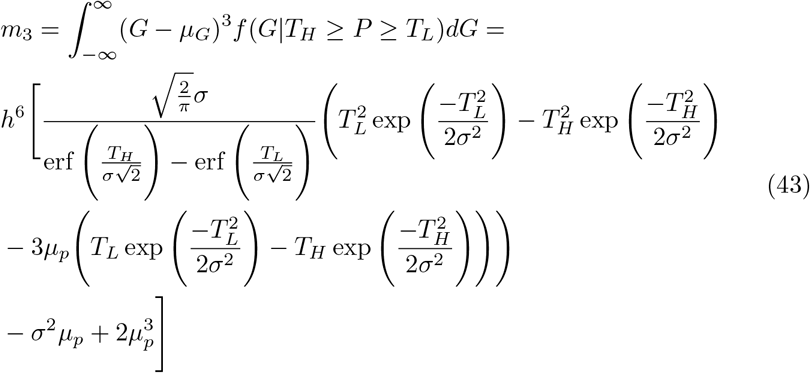

In the special case of directional selection above a threshold *T*

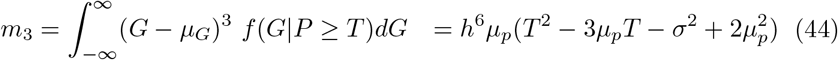

The traditional Fisher-Pearson skew can then be obtained as

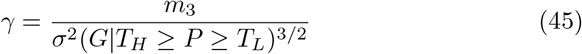

Similar to the mean, the skew for stabilizing and disruptive selection is zero. Comparing the calculated third moment for the breeding distribution above to the third moment of the phenotypic distribution post-selection gives us the relation *m*_3_ = *h*^6^*m*_3_(*P*).

### 8.2 Excess kurtosis

The final moment derived in this paper is the fourth moment which corresponds to the excess kurtosis of the breeding value distribution. Before selection, the kurtosis of the breeding value distribution is 3*h*^4^*σ*^4^ and excess kurtosis, the kurtosis differing from this value, is zero. Applying interval selection we find this changes

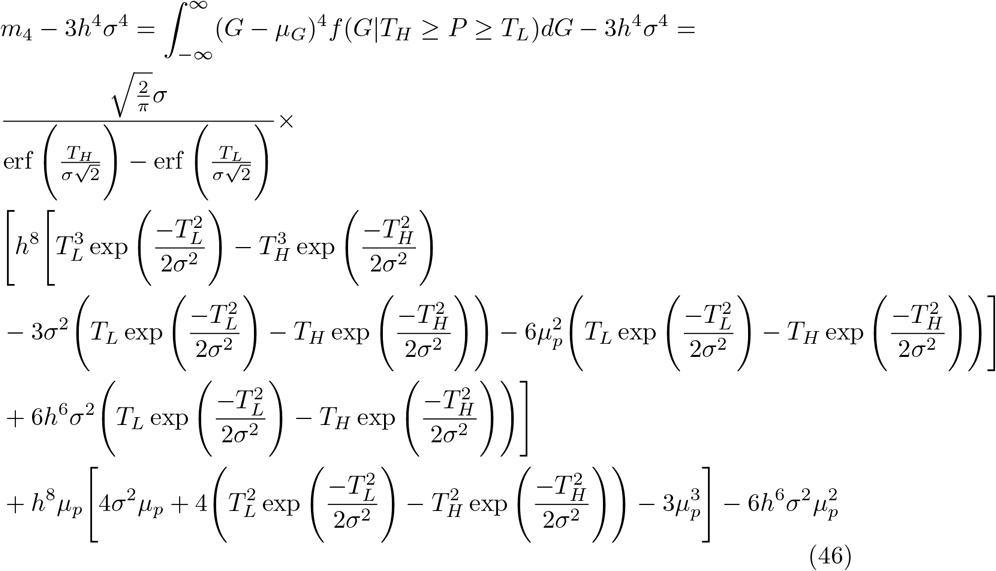

For stabilizing selection this reduces to

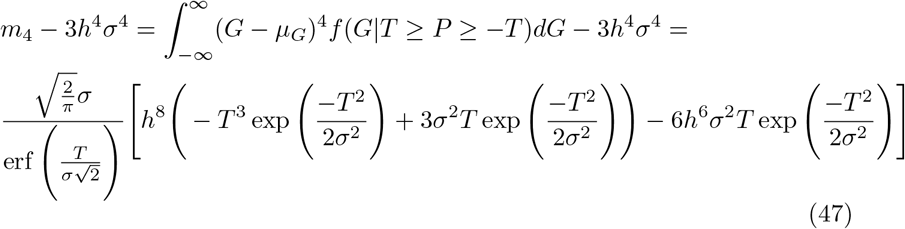

The final important selection we will deal with using the infinite population model will be the excess kurtosis of disruptive selection.

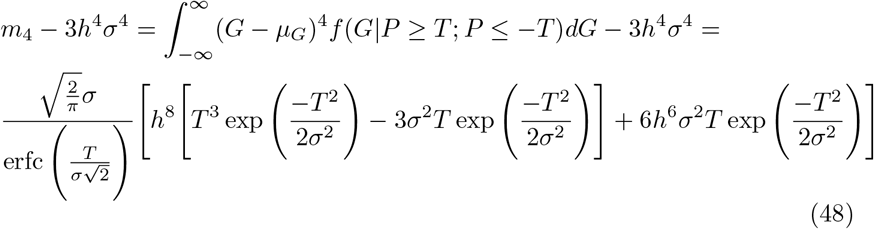

A key finding of the simulations in [Barton & Turelli, 1994] was that strong disruptive selection increases excess kurtosis in a large manner. We demonstrate that here showing the expression for disruptive selection. This result is confirmed as excess kurtosis sharply increases with the disruptive selection threshold in a way stabilizing and interval selection do not. In Figure 2 the graph of excess kurtosis versus disruptive selection threshold in standard deviations is demonstrated.

**Fig. 2.**
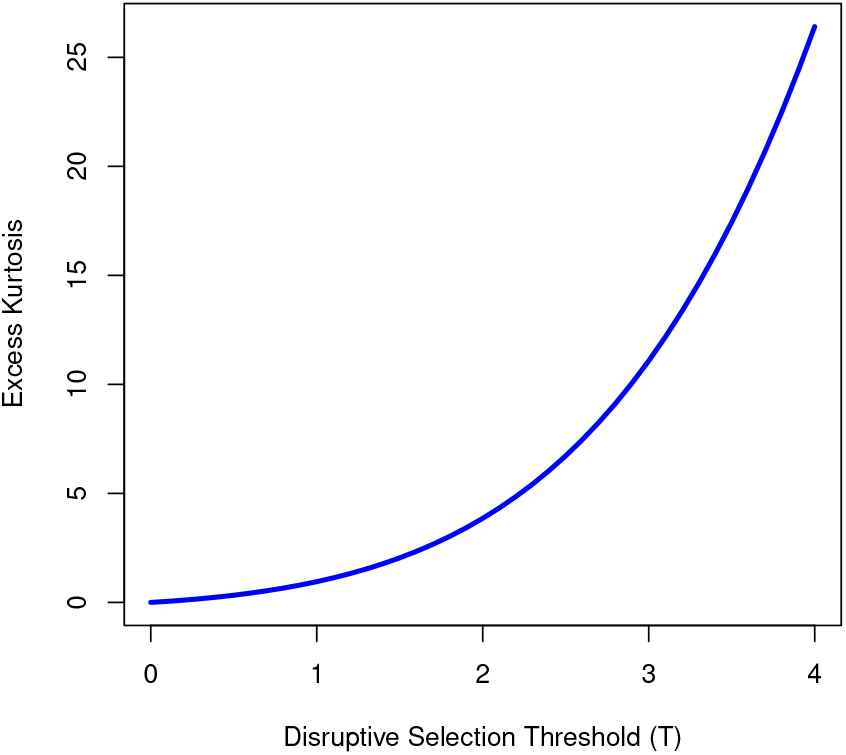
The excess kurtosis in the selected parent population generated by disruptive selection based on the threshold of selection *T* standard deviations. Excess kurtosis is reduced to 1/8 this value in the progeny by recombination. Created in R.

#### 8.2.1 Excess kurtosis of truncated directional selection

The equation for truncated directional selection is omitted here due to factors which make its interpretation problematic in comparison to real populations or simulated populations of finite size. The kurtosis, being the fourth moment, is rapidly inflated by only a few small outliers given its fourth power exponent and this causes problems of interpretation between a theoretical calculation based on infinite population size, shown here, and a finite population where very few members exist more than thee or four standard deviations from the mean. Based on an infinite population, the theoretical calculation of excess kurtosis in truncated directional selection is largely inflated as the selection threshold moves away from the mean where the theoretical and calculated values can be an order of magnitude different due to the effect of probabilistic rare values in the theoretical model. Simulation shows that the excess kurtosis of truncated directional selection, as well as interval selection for intervals far from the mean, suffers from bias due to the absence of large outliers in simulated or real data. Therefore, calculated values from simulation on realistically sized populations offers a superior measure of excess kurtosis in most cases of selection where the selected population mean is far from the original population mean. The stabilizing selection and disruptive selection equations tend to give accurate results since the mean of the post-selection population is unchanged and extreme values on both sides of the distribution balance each other out.

#### 8.2.2 Breakdown of higher moments by recombination

Barton and Turelli [Barton & Turelli, 1991, Barton & Turelli, 1994] discussed the third and fourth moments in terms of linkage disequilibrium amongst three and four loci similar to the pairwise linkage disequilibrium demonstrated by the Bulmer Effect. Consistent with their results, higher order linkage disequilibriums are broken down with recombination during meiosis. The basic pattern of recombination probabilities was described in detail by two papers by Geiringer [Geiringer, 1944, Geiringer, 1945]. In short, when all *m* loci are unlinked, and the so-called linkage distribution has a uniform distribution of values (1*/*2)^*m*^, the recombination probabilities are 2(1*/*2)^*m*^ = (1*/*2)^*m*−1^. Thus for skew, the progeny value is 1/4 of the selected parent value while excess kurtosis is reduced to 1/8 of the selected value. In other words, both higher moments are rapidly reduced by recombination in just one to two generations.

## 9 Discussion

The statistical moments of quantitative traits in a population undergoing selection have long been researched but previously, only linear regression, indirect, or computational tools were available. Primarily using recently available mathematical tables, the exact expressions for these moments have been derived for the first time confirming what was long known about the first and second moments but proving what was suspected about higher moments, particularly the fourth moment in disruptive selection scenarios. In short, departures for normality induced by selection are small and short-lived for the skew and excess kurtosis under most scenarios. The large values of these moments in selected parents is tempered by their reduction through recombination to the progeny generation. By the third generation, even large values of excess kurtosis are reduced to 1/64 of their original value.

The explicit derivation of the probability density function for breeding values post-selection has allowed other previously unknown insights. One is the ability to mathematically express the effect of recombination on the breeding values between generations. While the expression is only fully accurate for two loci linkage disequilibrium, the distortion of the population variance, it still allows a substantial investigation on how recombination shapes the breeding value distribution through generations.

Finally, far from just determining the Bulmer Effect, it was demonstrated how the change in phenotypic variance from selection can explain changes in the GXE covariance and the phenotype-breeding value covariance amongst selected parents. This could possibly have applications, for example in agriculture, where one wants to determine what environmental variables are important in determining the phenotype of the population in a selected sample.

Quantitative genetics is an ever evolving topic an even old and “basic” models can provide new insights or starting points for analysis. It is hoped that the results here will allow more advanced analytical methods to be applied to understand the effects of the evolution of quantitative traits.

## 10 Declarations

Funding: Not applicable Conflicts of interest/Competing interests: Not applicable Availability of data and material: Not applicable Code availability: Not applicable

## A Key integrals of the error function

The error function is defined as erf = *∫* exp (−*x*^2^)*dx*, but more generally

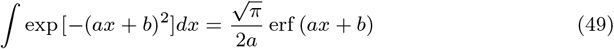

Section 1.1.1 Equation 1 [Korotkov & Korotkov, 2020]

The general integral of erf itself

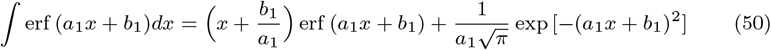

Equation 1.9.1 [Korotkov & Korotkov, 2020]

The definite integral of the product of the erf and the basic exponential defining density in a Gaussian over the domain [−*∞, ∞*]

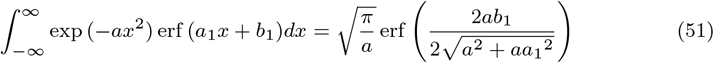

Section 2.7.1 Equation 6 [Korotkov & Korotkov, 2020]

The next group of integrals are used to calculate moments for the distributions of the form exp (−*ax*^2^) erf (*a*_1_*x* + *b*_1_)

First moment

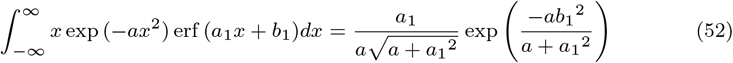

Section 2.7.2 Equation 4 [Korotkov & Korotkov, 2020]

Second moment

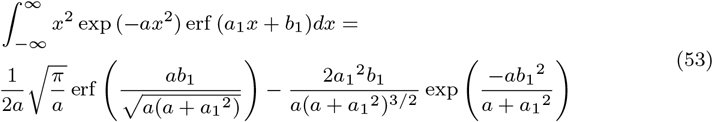

Section 2.7.3 Equation 9 (*n* = 2)[Korotkov & Korotkov, 2020]

Third moment

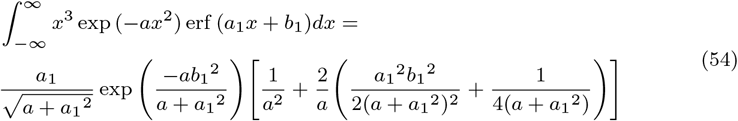

Section 2.6.2 Equation 4 (*m* = 1) [Korotkov & Korotkov, 2020]

Fourth moment

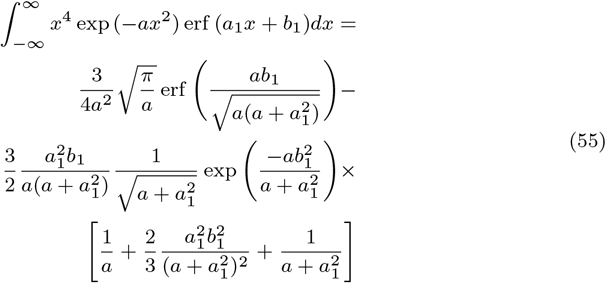

Section 2.7.3 Equation 9 (*n* = 4) [Korotkov & Korotkov, 2020]

## Notes

### Competing Interest Statement

The authors have declared no competing interest.

